# Exonic splice variant discovery using in vitro models of inherited retinal disease

**DOI:** 10.1101/2024.05.06.592740

**Authors:** Nathaniel Kevin Mullin, Laura R. Bohrer, Kristin R. Anfinson, Jeaneen L. Andorf, Robert F. Mullins, Budd A. Tucker, Edwin M. Stone

**Author notes:** **Corresponding author:** Edwin M. Stone, M.D., Ph.D., 375 Newton Road, 4111 MERF, The University of Iowa, Iowa City, IA 52242.

## Abstract

Correct identification of the molecular consequences of pathogenic genetic variants is essential to the development of allele-specific therapies. However, in some cases, such molecular effects may be ambiguous following genetic sequence analysis alone. One such case is exonic, codon-altering variants that are also predicted to disrupt normal RNA splicing. Here, we identify such cases in the context of inherited retinal disease. *NR2E3* c.932G>A (p.Arg311Gln) is a variant commonly associated with Enhanced S Cone Syndrome (ESCS). Previous studies using mutagenized cDNA constructs have shown that the arginine to glutamine substitution at position 311 of NR2E3 does not meaningfully diminish function of the rod-specific transcription factor. Using retinal organoids, we explored the molecular consequences of *NR2E3* c.932G>A when expressed endogenously during human rod photoreceptor cell development. Retinal organoids carrying the *NR2E3* c.932G>A allele expressed a transcript containing a 186-nucleotide deletion of exon 6 within the ligand binding domain. This short transcript was not detected in control organoids or control human donor retina samples. A minigene containing exons 5 and 6 of *NR2E3* showed sufficiency of the c.932G>A variant to cause the observed splicing defect. These results support the hypothesis that the pathogenic *NR2E3* c.932G>A variant leads to photoreceptor disease by causing a splice defect and not through an amino acid substitution as previously supposed. They also explain the relatively mild effect of Arg311Gln on NR2E3 function *in vitro*. We also used *in silico* prediction tools to show that similar changes are likely to affect other inherited retinal disease variants in genes such as *CEP290, ABCA4*, and *BEST1*.

## INTRODUCTION

Enhanced S-cone syndrome (ESCS) (OMIM 268100) is an inherited retinal disease characterized by developmental malformation of the outer neural retina ^1-5^. Classic features of this disease include night blindness and increased sensitivity to blue light ^1^. Loss of function of the rod-specific transcription factor nuclear receptor subfamily 2, group E, member 3 (NR2E3, HGNC:7974) is the most common cause of ESCS ^1^. While for many patients ESCS is a relatively stationary disorder ^6^, NR2E3 variants can cause a more severe form of retinal degeneration known as Goldmann-Favre Syndrome. Interestingly, a clear genotype-phenotype correlation has yet to be identified ^7^.

More than 80 pathogenic variants in *NR2E3* have been described to date ^8^. One of the more common ESCS-causing mutations in *NR2E3* is a coding sequence variant at position 932 (NM_014249.4:c.932G>A), which converts codon 311 from C<u>G</u>G to C<u>A</u>G ^1^. This variant has been previously thought to exert its pathogenic effect through a resulting arginine to glutamine residue change at this position (p.Arg311Gln). Residue 311 is found within the ligand-binding domain of NR2E3, further supporting the potential for protein dysfunction following normal amino acid sequence disruption. However, conflicting evidence from *in vitro* experiments and *in silico* modeling exists for the p.Arg311Gln substitution being sufficient for loss of NR2E3 protein function ^9-11^. Notably, expression of mutangenized *NR2E3* cDNAs to generate protein containing the Arg311Gln change have not shown markedly diminished NR2E3 behavior in terms of co-factor binding or ability to act at known regulatory elements ^9-11^.

To evaluate pathobiology of genetic variants such as p.Arg311Gln, use of disease relevant tissue types that natively express the gene of interest are more likely to provide meaningful results. Unlike the skin, which can readily be biopsied for direct evaluation, surgical sampling of the neural retina would result in loss of visual function in the sampled area and as such not something that is done unless absolutely necessary (e.g., to identify a pathogen that would otherwise destroy the retina). Patient iPSC-derived retinal organoids, which follow normal retinal developmental programs, express retina-specific genes ^12-14^, and replicate retina-specific isoform expression ^12,15,16^, are the ideal alternative to primary retinal tissue for evaluation of the molecular consequences of pathogenic variants. In this study, we investigated the molecular significance of the *NR2E3* c.932G>A (p.R311Q) variant using retinal organoids. We found that in human retinal cells, c.932G>A creates an alternative 3’ splice site within exon 6. The resulting transcript lacks 186 bases of the 5’ end of exon 6, presumably causing an in-frame deletion of 62 amino acids from NR2E3’s ligand binding domain. We showed that this change was predictable based on *in silico* models of variant effect. We also found that in a cohort of 1,000 inherited retinal disease families, only 4 other single nucleotide variants currently annotated as missense variants reach the same prediction confidence threshold for splice site alteration. Using cultured fibroblasts from a patient carrying one such variant in the gene encoding centrosomal protein 290 (*CEP290*, HGNC:29021; c.1711G>A (p.Gly571Arg)), we confirmed that this variant leads to expression of a longer isoform containing a frame-shifting intronic inclusion. Together, these results shed new light on the pathogenic mechanism of a common pathogenic *NR2E3* variant and highlight a potential gap in the annotation of exonic splice variants in inherited retinal disease genes.

## METHODS

### *In silico* variant effect prediction

Twenty-one *NR2E3* variants were retrieved from ClinVar (https://www.ncbi.nlm.nih.gov/clinvar/, accessed December 2023). Only variants classified as having a clinical significance of *“Pathogenic”* or “*Likely Pathogenic”* and a molecular consequence of “*Missense”* were used. SpliceAI (Version 1.3.1) was run against the hg19 human reference genome assembly with default parameters. For AlphaMissense analysis, pathogenicity scores for variants of interest were retrieved from the genome-wide results previously generated by Google DeepMind (AlphaMissense_aa_substitutions.tsv, accessed December 2023). Results from these analyses were processed and plotted using R (version 4.3.0).

### Patient-derived iPSCs

This study was approved by the Institutional Review Board of the University of Iowa (project approval #200202022) and adhered to the tenets set forth in the Declaration of Helsinki. Patient iPSCs were generated previously from two individuals with molecularly confirmed enhanced S cone syndrome (ESCS) as described ^17^.

### Retinal organoid differentiation

Retinal differentiation was performed as described previously with minor modifications ^17,18^. Briefly, iPSCs were cultured on laminin 521 coated plates in E8 medium. Embryoid bodies (EBs) were lifted with ReLeSR (STEMCELL Technologies) and transitioned from E8 to neural induction medium (NIM-DMEM/F12 (1:1), 1% N2 supplement, 1% non-essential amino acids, 1% Glutamax (Thermo Fisher Scientific), 2 μg/mL heparin (Sigma-Aldrich) and Primocin (InvivoGen)) over a four-day period. On day 6, NIM was supplemented with 1.5 nM BMP4 (R&D Systems). On day 7, EBs were adhered to Matrigel coated plates (Corning). BMP4 was gradually transitioned out of the NIM over seven days. On day 16, the media was changed to retinal differentiation medium (RDM - DMEM/F12 (3:1), 2% B27 supplement (Thermo Fisher Scientific), 1% non-essential amino acids, 1% Glutamax and 0.2% Primocin). On day 25-30 the entire EB outgrowth was mechanically lifted using a cell scraper and transferred to ultra-low attachment flasks in 3D-RDM (RDM plus 10% fetal bovine serum (FBS)); Thermo Fisher Scientific), 100 μM taurine (Sigma-Aldrich), 1:1000 chemically defined lipid concentrate (Thermo Fisher Scientific), and 1 μM all-trans retinoic acid (until day 100; Sigma-Aldrich). The cells were fed three times per week with 3D-RDM until the point of RNA isolation.

### RNA isolation and cDNA synthesis

RNA was isolated using the NucleoSpin RNA Kit (Machery-Nagel) following the manufacturer’s protocol. In addition to the on-column DNaseI treatment, an additional DNA digestion was performed with RQ1 RNase-Free DNase following the manufacturer’s protocol for RT-PCR (Promega). cDNA synthesis was carried out using the Superscript IV VILO system (Invitrogen) following the manufacturer’s protocol. RNA was stored at -80°C in water and cDNA was stored at - 20°C.

### RT-PCR

Custom oligos were designed using the PrimerQuest online tool (Integrated DNA Technologies) and synthesized by Integrated DNA Technologies. Oligos were resuspended to a concentration of 100µM in water and stored at -20°C. The polymerase chain reactions (PCRs) were carried out using BIOLASE DNA polymerase (Bioline) with 35 cycles, an annealing temperature of 52°C for *NR2E3* and 50°C for *CEP290*, and a 30 second extension at 72°C. PCR products were electrophoresed on 2% agarose gels with SYBR Safe stain (Invitrogen).

### RT-PCR Amplicon Sequencing

Bands of interest were cut from the gels, and the amplicons were purified using the QIAquick Gel Extraction kit following manufacturer’s protocol (Qiagen). Sanger sequencing was performed by Functional Biosciences (Madison, Wisconsin). The resulting chromatograms were visualized using SnapGene (version 6.1.2).

### Long-read gDNA amplicon sequencing

The entire ∼8kb *NR2E3* locus was amplified using *NR2E3* F and *NR2E3* R primers (**Table 1**) with the Expand Long Template PCR System (Roche) buffer system 1. PCR was performed for 30 cycles with an annealing temperature of 58°C and an extension of 11 minutes with extension time increasing 5 seconds per cycle for the last 20 cycles, following the manufacturer’s protocol. PCR products were prepared for long-read sequencing using the Native Barcoding Kit 24 V14 (Oxford Nanopore Technologies) following the manufacturer’s protocol. Briefly, sample index barcodes and sequencing adapters were added to amplicons via ligation and libraries were pooled for sequencing on an R10.4.1 flow cell on the MinION Mk1B instrument (Oxford Nanopore Technologies). Base calling and demultiplexing was performed with Guppy (version 6.3.8) using the dna_r10.4.1_e8.2_260bps_sup.cfg configuration and reads were mapped to the hg38 reference genome with Minimap2 (version 2.24) ^19^. Aligned reads were visualized with the Integrative Genomics Viewer desktop application (version 2.16.2).

**Table 1.**
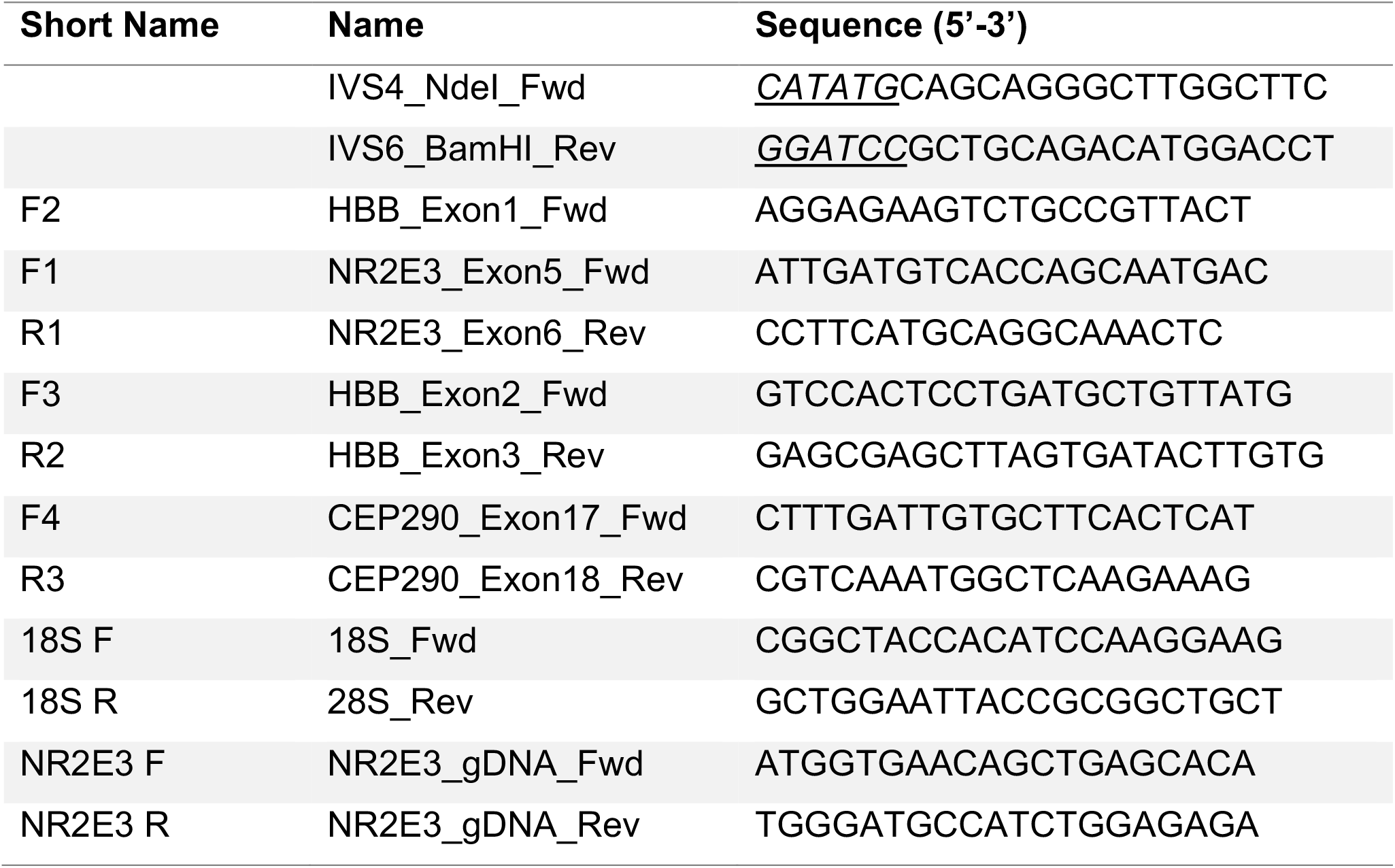
Primers used for RT-PCR, dPCR, and minigene construction.

**Table 2.** Cell lines used.

| Short Name | Sample | Differentiation Timepoint | *NR2E3* Genotype | *CEP290* Genotype |
| --- | --- | --- | --- | --- |
| P1 | Patient 1 | D83 | c.932G>A<br>(p.R311Q)/<br>c.1192-A>C | n/a |
| P2 | Patient 2 | D105 | c.932G>A<br>(p.R311Q)/<br>c.219G>C<br>(p.R73S) | n/a |
| P3 | Patient 3 | n/a | n/a | c.1711G>A (p.Gly571Arg)/<br>c.7048C>T (p.Gln2350*) |
| Ctl | Control 1 | D161 | n/a | n/a |
| D1 | Donor 1 | n/a | n/a | n/a |
| D2 | Donor 2 | n/a | n/a | n/a |
| D3 | Donor 3 | n/a | n/a | n/a |
| iPS | iPSC Control | n/a | n/a | n/a |

### Long-read cDNA sequencing

Long-read sequencing of *NR2E3* cDNA was carried out using the MinION sequencing platform (Oxford Nanopore Technologies). cDNA was generated using the Chromium Next GEM Single Cell 3’ Kit v3.1 (10X Genomics) from dissociated retinal organoids as described previously ^5^. Libraries were generated from this cDNA using the PCR-cDNA Sequencing Kit (SQK-PCS111) and sequenced with a R9.4.1 flow cell on the MinION Mk1B instrument (Oxford Nanopore Technologies). Reads were processed with Guppy and Minimap2 as described above.

### Minigene construction

The region containing exon 6 of *NR2E3* (i.e., IVS5-IVS6) was amplified from genomic DNA isolated from Patient 1 using Q5 High-Fidelity DNA Polymerase (New England Biolabs) with 35 cycles, an annealing temperature of 71°C, and 1 minute extension at 72°C. Primers containing *NdeI* and *BamHI* restriction sites were used (**Table 1**). The resulting amplicon was cloned into the pCR2.1-Blunt-II backbone using the Zero Blunt TOPO PCR Cloning Kit following manufacturer’s protocol (Invitrogen). Clones were screened by Sanger sequencing to identify clones containing either G or A at c.932. Plasmids from selected clones were digested with *NdeI* and *BamHI* (New England Biolabs) and the resulting insert gel purified as above and ligated into the modified pcAT7-Glo1 vector (a gift from Brenton Graveley, Addgene Plasmid # 160996) prepared with *NdeI* and *BglII* digest and gel purification. Minipreps of the resulting clones were prepared using the NucleoSpin Plasmid kit following the manufacturer’s protocol (Machery Nagel) and confirmed by diagnostic *EcoRI* digest and Sanger sequencing across the insert.

### Minigene transfection

HEK293T cells (ATCC) were cultured in MEM α (Gibco) with 10% fetal bovine serum (Atlas Biologicals) and 100µg/mL Primocin (InvivoGen). Cells were transfected at ∼60% confluence with Lipofectamine 3000 following the manufacturer’s protocol. 500ng of minigene construct was used per well of a 24-well plate. As a transfection control, a copGFP-expressing plasmid was used. 48 hours following transfection, total RNA was isolated using the NucleoSpin RNA kit (Machery Nagel) as described above.

## RESULTS

### Predicted molecular consequences of genetic variants in *NR2E3*

RNA splicing is initiated via conserved motifs that define splice junctions. Models built on these patterns offer a predictive tool for splice site prediction ^20^. Similarly, predictive models of the consequences of amino acid changes on protein structure and function have been constructed ^21^. We used SpliceAI and AlphaMissense to ask whether the *NR2E3* c.932G>A (p.Arg311Gln) variant and other pathogenic missense variants within *NR2E3* would be predicted to cause changes in RNA splicing or protein function (**Figure 1**). Twenty-one missense variants reported in ClinVar to be either Pathogenic or Likely Pathogenic were used in this analysis. For each variant, the AlphaMissense score and SpliceAI Delta Scores were plotted. *NR2E3* p.R311Q returned the lowest AlphaMissense score of this collection (0.093) as well as the highest Delta Score (for Acceptor Gain) from SpliceAI (0.69). No other pathogenic missense variant in *NR2E3* was predicted as strongly to alter splicing.

**Figure 1.**
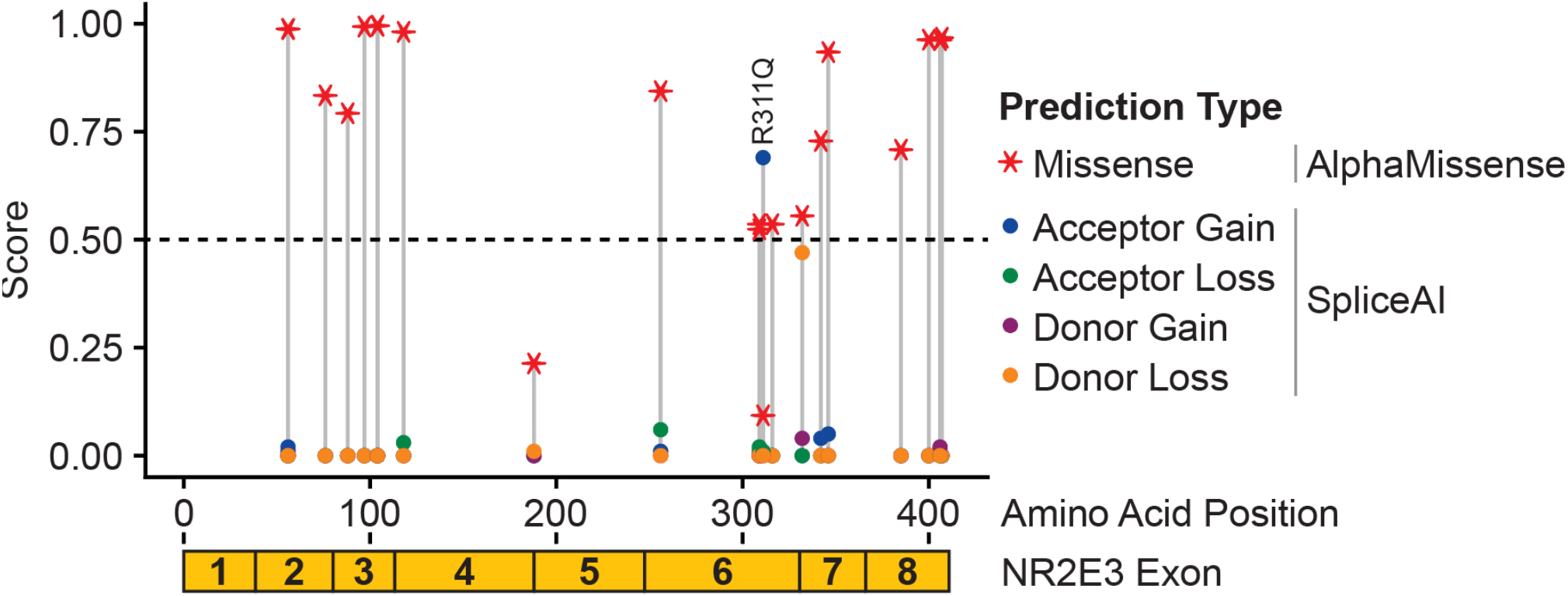
Missense variant effect prediction in NR2E3. For 21 missense variants classified as Pathogenic or Likely Pathogenic, AlphaMissense and SpliceAI prediction scores are shown. Variants are shown based on position with the NR2E3 coding sequence. Each variant is represented by a vertical gray line and colored points along the line represent the magnitude of prediction strength for missense or splicing impact. The p.R311Q variant is labeled and exhibits a higher score for splice acceptor gain (blue circle) and a lower score for missense impact (red star).

### Truncated NR2E3 transcript expression in c.932G>A organoids

We next tested the effects of c.932G>A on NR2E3 splicing directly using patient-derived retinal cells carrying this variant. cDNA derived from ESCS patient retinal organoids was sequenced using a long reads and splice junctions were counted (**Figure 2A, B**). The patient carried a previously described compound heterozygous *NR2E3* genotype composed of variants c.219G>C and c.932G>A (**Figure 2**) ^17^. Out of a total of 58, 28 reads exhibited a splicing between the 3’ end of exon 5 and the middle of exon 6 at the location of the c.932G>A variant (**Figure 2B**). No other aberrant splicing was observed, implying that c.932G>A creates an alternative splice acceptor within exon 6.

**Figure 2.**
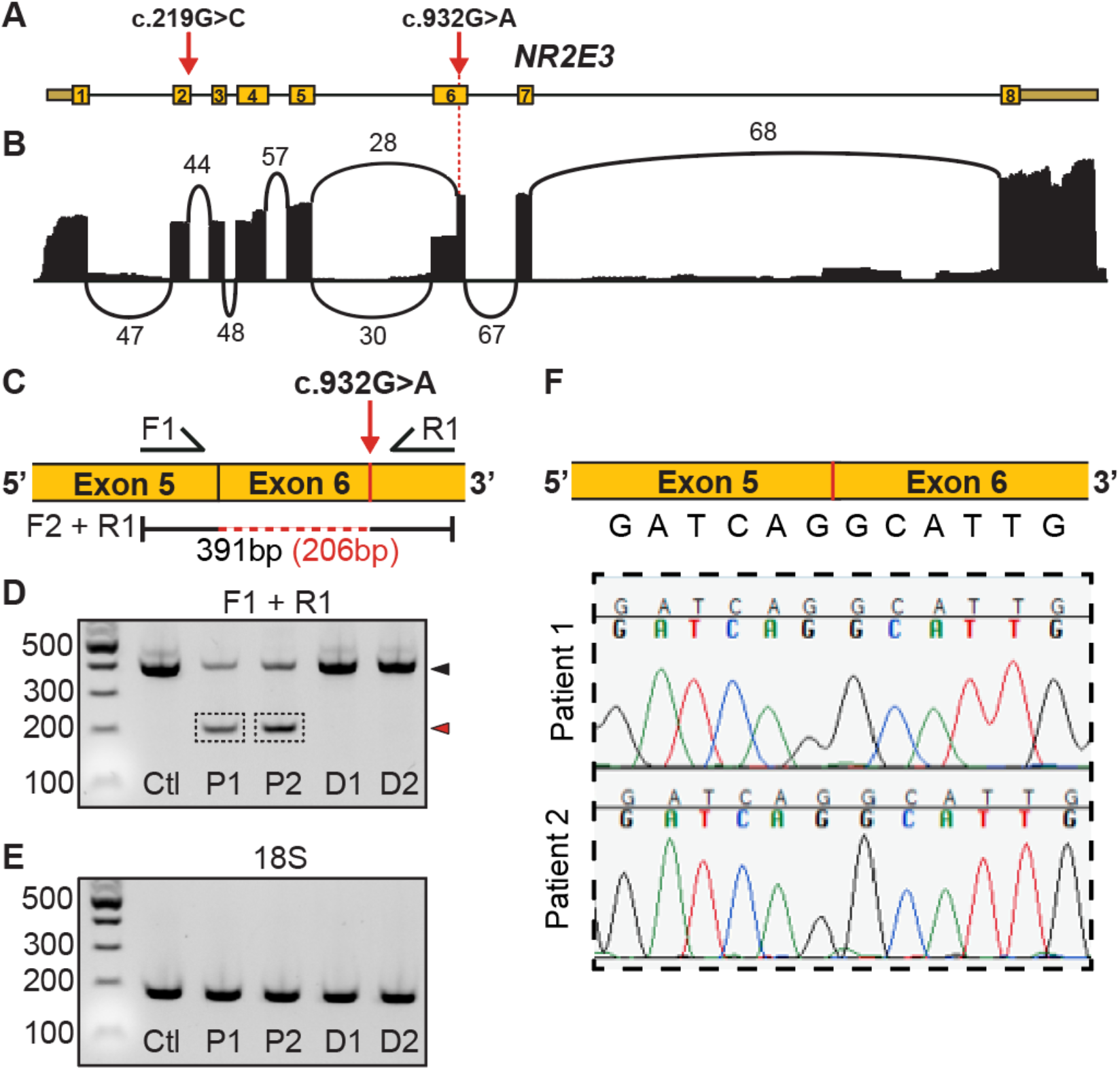
Retinal organoid expression of a misspliced transcript near the c.932G>A variant. **A)** Canonical NR2E3 gene structure including 8 exons. The positions of two pathogenic variants in exons 2 and 6 (c.219G>C and c.932G>A respectively) are shown. **B)** Sashimi plot representing long-read RNA sequencing reads mapping to *NR2E3*. The number of reads spanning exons is shown. 67 reads span from exon 6 to exon 7. 30 reads span from exon 5 to the canonical splice acceptor at the 5’ end of exon 6. 28 reads span from exon 5 to the presumptive novel splice acceptor in exon 6. **C)** PCR primer design to span the truncated portion of exon 6 in misspliced transcripts. Amplicon originating from normal transcript is expected to be 391bp while that from misspliced transcript is expected to be 206bp. **D)** RT-PCR using F1 and R1 primers shown in **C**. In control organoids (Ctl), only full-length amplicon is observed (black arrowhead). In organoids from two unrelated patients with compound heterozygous NR2E3 genotypes (P1, P2), both full length and truncated amplicon (red arrowhead) is observed. Two unrelated human donor retina samples (D1, D2) express only properly spliced NR2E3 transcript. **E)** RT-PCR for 18S rRNA serves as a loading control. **F)** Sanger sequencing of the shorter amplicon from P1 and P2 (shown boxed in **D**) shows consistent missplicing of NR2E3 exon 5 into the predicted alternative splice acceptor within exon 6.

To further confirm allele-specific missplicing of *NR2E3* transcript, primers spanning exons 5 and 6 of *NR2E3* were designed and used for RT-PCR of RNA from D80-D100 retinal organoids (**Figure 2C**, see **Table 3** for sample information). Two lines from unrelated ESCS patients were used. RT-PCR from both patient lines yielded two amplicons; one of the expected size for canonically spliced *NR2E3* and a smaller amplicon showing the expected size following splicing using an exonic splice site created at c.932G>A (**Figure 2D, E**). To confirm the identity of these amplicons, The smaller amplicon from each sample was isolated and sequenced. Both patient lines showed the same splice junction of the 3’ end of exon 5 being spliced to an alternative acceptor within exon 6 at the expected position based on RNA-seq data (**Figure 2F**).

The presence of normal exonic and intronic genomic DNA sequence in the two patients was confirmed by long-read sequencing of an amplicon spanning the entire *NR2E3* locus (**Figure S1A, B**). Both patients exhibit compound heterozygous alleles. Other unique benign variants were observed in both samples. No structural variants (such as deletion involving exon 6) were observed, further supporting c.932G>A as being causal for transcript missplicing.

### Minigene validation of c.932G>A sufficiency for exonic splice acceptor creation

To test the sufficiency of the c.932G>A variant for causing missplicing between exons 5 and 6 of *NR2E3*, we next constructed a minigene plasmid to express these exons in a non-retinal cell line. The sequence containing these exons (i.e., spanning *NR2E3* introns 4-6) was amplified and cloned into the pcAT7-Glo1 ^22^ vector between human hemoglobin subunit beta (*HBB*) exons 1 and 2 (**Figure 3A**). The resulting construct drives constitutive expression under the chimeric CAG promoter ^22,23^. Constructs containing both the reference (G) and alternate (A) base at c.932 were generated. Minigene constructs were transfected into HEK293T cells and RT-PCR using intron-spanning primers was used to amplify spliced template (**Figure 3B**) across various junctions. As in the organoid samples, a truncated cDNA was observed in cells transfected with the c.932G variant-containing minigene (**Figure 3C, D**) but not in the wildtype condition. No endogenous *NR2E3* expression in HEK293T cells was detected, consistent with previous reports of the retinal specificity of the gene. Notably, a small amount of normally spliced *NR2E3* fragment was observed in cells transfected with the variant-containing construct (**Figure 3C, D** black arrowhead). Normal splicing of *HBB* exons 2 and 3 was observed, indicating normal splicing of the expressed pre-mRNA in HEK293T cells (**Figure 3E**). 18S was used as a loading control (**Figure 3F**). Together, the results of this minigene experiment show that that *NR2E3* c.932G>A is sufficient to cause missplicing in a minimal *in vitro* system. They also demonstrate that such missplicing is not photoreceptor cell specific.

**Figure 3.**
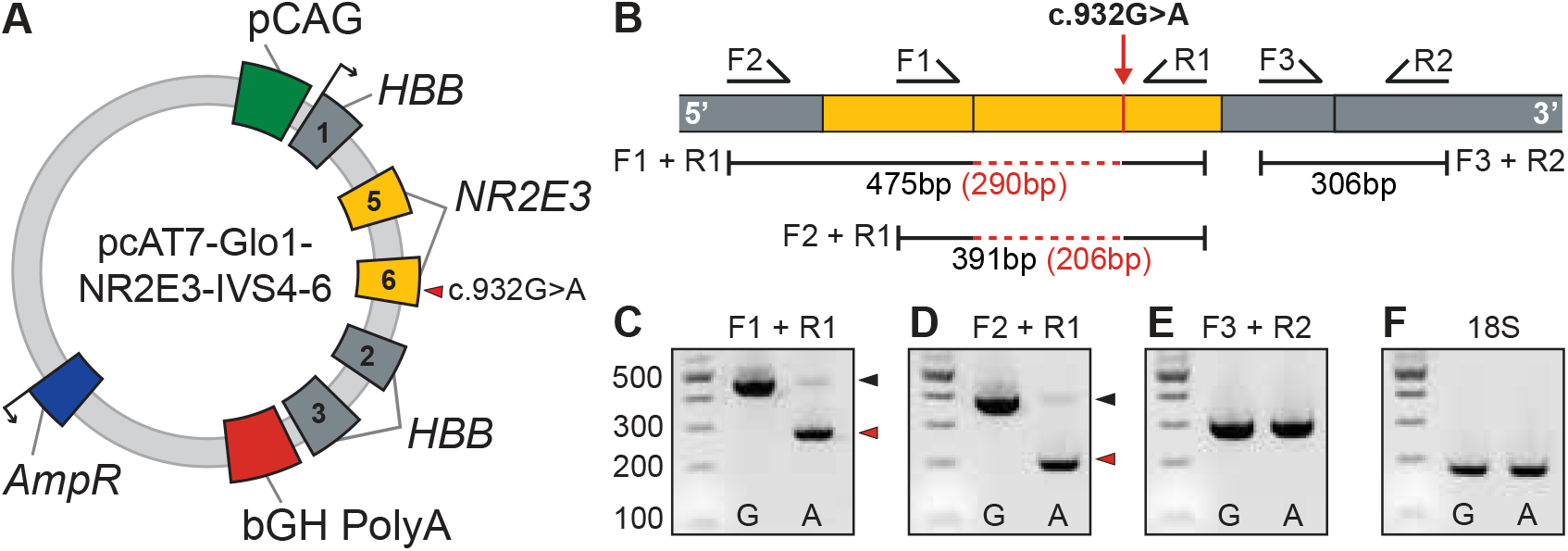
A NR2E3 minigene construct shows sufficiency of c.932G>A to cause missplicing. **A)** Plasmid design for testing splicing of NR2E3 exons 5 and 6 containing the c.932G>A variant. The NR2E3 locus was cloned into the pcAT7-Glo1 expression construct between HBB exons 1 and 2 and expressed in HEK 293T cells. **B)** Design of RT-PCR primers to assess splicing of minigene transcript. Splicing between HBB exons 2 and 3 serve as a control of normal splicing in this experimental system. **C, D)** Amplicons from samples transfected with the mutant (A) minigenes were 185 base pairs smaller than those from wildtype (G) constructs. Normal amplicons shown with black arrowheads, truncated amplicons shown with red arrowheads. Of note, a small amount of normal-length amplicon is seen in both mutant conditions (black arrowheads, “A” lanes). **E)** Normal splicing of HBB exons 2 and 3 is observed in both conditions. **F)** Expression of 18S RNA serves as a loading control.

### Predicting splicing consequences of inherited retinal disease missense variants

In a recent cohort of 1,000 inherited retinal disease families ^24^, 767 disease-causing alleles were identified. Of these, 529 (69%) involved a single nucleotide change. A high proportion of genes that cause inherited retinal disease are expressed only in the retina ^12^ or are spliced in a tissue-specific manner ^16^. Following our observation that *NR2E3* c.932G>A acts through an unexpected pathogenic mechanism, we next asked how common such a finding might be within this inherited retinal disease cohort. A total of 529 single-nucleotide variants were analyzed using SpliceAI and delta scores for gain or loss of splice sites in the alternate allele were returned (**Figure 4A**). Only 9/529 variants (1.7%) had a delta score at least as high as that of *NR2E3* c.932G>A. 14/529 variants (2.6%) had a score of at least 0.5, a threshold for confident cryptic splice site identification using SpliceAI ^20^. Of the top 9 variants, 5 had the highest delta score for Donor Gain, 3 for Donor Loss and 1 for Acceptor Gain (*NR2E3* c.932G>A). The variants were distributed among 7 genes, with 3 found in *ABCA4* and 1 found in each of six additional genes: *BEST1, CEP290, NPHP1, NR2E3, RHO*, and *USH2A*. Nonsynonymous, non-truncating variants from the set of 14 with a SpliceAI delta score greater than 0.5 were next analyzed with AlphaMissense (**Figure 4B**) to determine the predicted impact of the annotated amino acid substitution. 4/9 variants were predicted to be likely benign as missense variants while the other 5 were ambiguous. None of the 9 variants were predicted to be likely pathogenic as missense.

**Figure 4.**
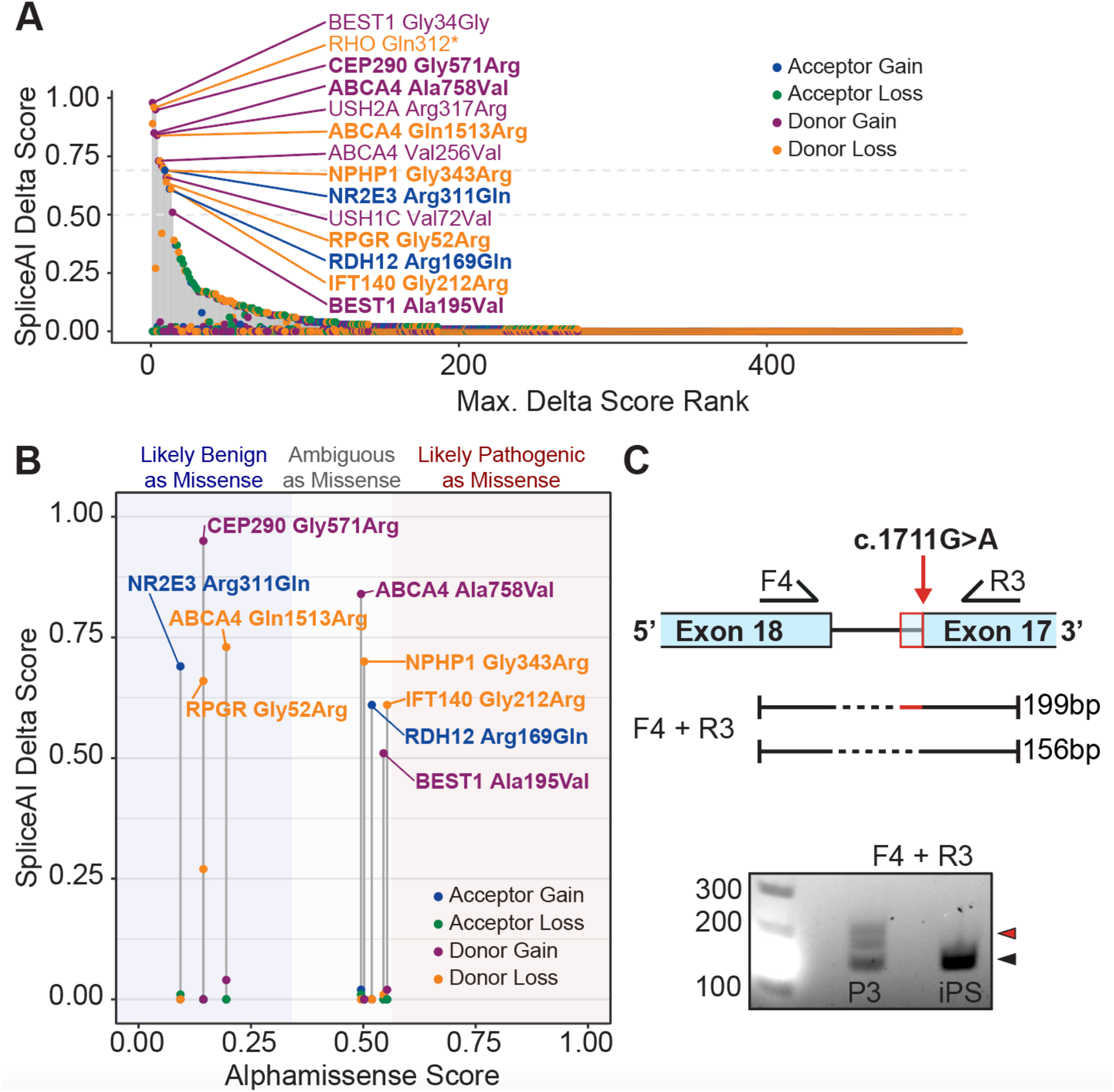
Splice variant prediction in an inherited retinal disease cohort. **A)** 529 variants were analyzed with SpliceAI, returning four delta scores per variant. Variants are shown in order of rank of maximum delta score. 9 variants (including *NR2E3* c.932G>A) had a score at least as high as *NR2E3* c.932G>A. 5 variants had a score lower than *NR2E3* c.932G>A but higher than 0.5, a common threshold for SpliceAI prediction. Most variants (515) had maximum delta scores lower than 0.5. **B)** Only non-synonymous and non-truncating variants with a maximum delta score of at least 0.5 from **A** are shown (9/14 variants from **A**). The gene symbol and predicted amino acid change is shown for each variant. On the x-axis, the AlphaMissense score is shown. **C)** RT-PCR was performed between *CEP290* exons 17 and 18 to assess missplicing caused by the c.1711G>A (p.Gly521Arg) variant, as predicted by the analysis in **B**. A larger product consistent with intron inclusion (red arrowhead) was observed in a compound heterozygous patient line (P3) but not a control line (iPS).

To further validate the above in silico predictions, we performed RT-PCR using RNA from a compound heterozygous patient fibroblast line carrying the *CEP290* c.1711G>A (p.Gly571Arg) variant (**Figure 4C**). Of the collection analyzed by SpliceAI and AlphaMissense, this variant was most highly predicted to alter splicing and was predicted to be likely benign as an amino acid substitution (**Figure 4B**). Indeed, dermal fibroblasts from a patient carrying this *CEP290* variant expressed a longer isoform of the transcript, presumably due to partial intron inclusion between exon 17 and 18. The *CEP290* c.1711G>A variant is predicted by SpliceAI to abolish the canonical splice donor of exon 17 and lead to preferential usage of a cryptic donor within the following intronic sequence. Together, this causes a frame-shifting intronic inclusion, as observed by RT-PCR (**Figure 4C**).

## DISCUSSION

Many variants that have been reported to cause inherited retinal disease are located within genes expressed exclusively in the retina. As it is difficult to obtain affected retinal tissue, the molecular consequences of pathogenic variants are often hard to evaluate. For this reason, the protein-level consequence of a genetic variant is often predicted based on codon disruption or known splice sites. Here, we show that *in vitro* models of inherited retinal disease such as patient iPSC-derived retinal organoids can be used to investigate variant effects and demonstrate that presumed missense variants actually cause missplicing of transcript when expressed.

Previous studies of the *NR2E3* c.932G>A (p.R311Q) variant have largely utilized mutangenized wildtype cDNA to compare variants using cell culture-based systems ^9-11^. While these strategies allow for highly controlled expression of different variants, they do not allow one to observe changes to splicing since only exonic sequences are present. In light of the findings presented here, the inability of c.932G>A to cause significant dysfunction of NR2E3 in previous studies can be explained by the splice modulating mechanism, indicating that in fact, an arginine to glutamine change at position 311 may not be pathogenic. Together, these results indicate that a modulation of splicing to express a full length *NR2E3* cDNA containing the c.932G>A variant may be sufficient to restore normal NR2E3 function. Further experiments must be performed to confirm this hypothesis. Additionally, since NR2E3 is a transcription factor active in early retinal development, it remains an open question whether restoring normal *NR2E3* expression in the adult retina would restore any visual function or prevent progressive retinal degeneration.

While nonsense or missense variants may require viral vector-mediated gene augmentation, splice variants may be amenable to modulation by antisense oligonucleotides. Previous work has shown that such treatment approaches can be successful in inherited retinal disease caused by errors in splicing of photoreceptor cell genes ^25^. Expanding the catalog of known splice variants to include variants currently annotated as missense variants (such as those in **Figure 4**) may permit new treatment strategies for several inherited retinal diseases.

In this study the *in-silico* splice prediction software SpliceAI was used to predict if known pathogenic missense mutations were likely to alter transcript splicing. In addition to c.932G>A (p.R311Q), missense variants in 13 genes previously reported ^24^ to be disease causing were identified. Using patient derived fibroblasts, we were able to demonstrate that the C.1711C>A (p.G571R) variant in CEP290 caused mis-splicing of the *CEP290* transcript, resulting in inclusion of a portion of intron 17 that was absent in the normal control (**Figure 4**). While these findings demonstrate the utility of SpliceAI, they also highlight the importance of using patient derived cells to validate *in-silico* predictions. Unlike *CEP290*, which is expressed in dermal fibroblasts, the remaining 12 variants identified are expressed in retina specific genes and as such require generation of patient iPSCs and derivation of retinal organoids. As this strategy is both labor and resource intensive it is most suitable for validation of variants whose pathogenicity are suspected but unknown. For instance, using patient derived retinal organoids we recently demonstrated that a deep intronic variant in the gene ABCA4 creates a cryptic splice site and miss-splicing in photoreceptor and RPE cells ^26^. In the latter study, data pertaining to the prevalence of the novel intronic variant in cases vs. controls allowed us to narrow the list of disease-causing candidates. One could envision using SpliceAI in a similar fashion; that is, as a hypothesis generating tool. Regardless, to support the molecular diagnosis of an unknown genetic variant, validation of *in-silico* predictions using relevant model systems such as patient derived retinal organoids is required.

In conclusion, this work shows that correct identification of a variant’s effect on a protein -- in this case, deletion of a large portion of its ligand-binding domain -- can be ascertained through studies based on disease modeling in human cells and assays at the transcript-level. Future work will be required to understand how the deletion caused by c.932G>A alters NR2E3’s ability to interact with co-factors and carry out its role in photoreceptor development.

## SUPPLEMENTAL FIGURE

**Figure S1.**
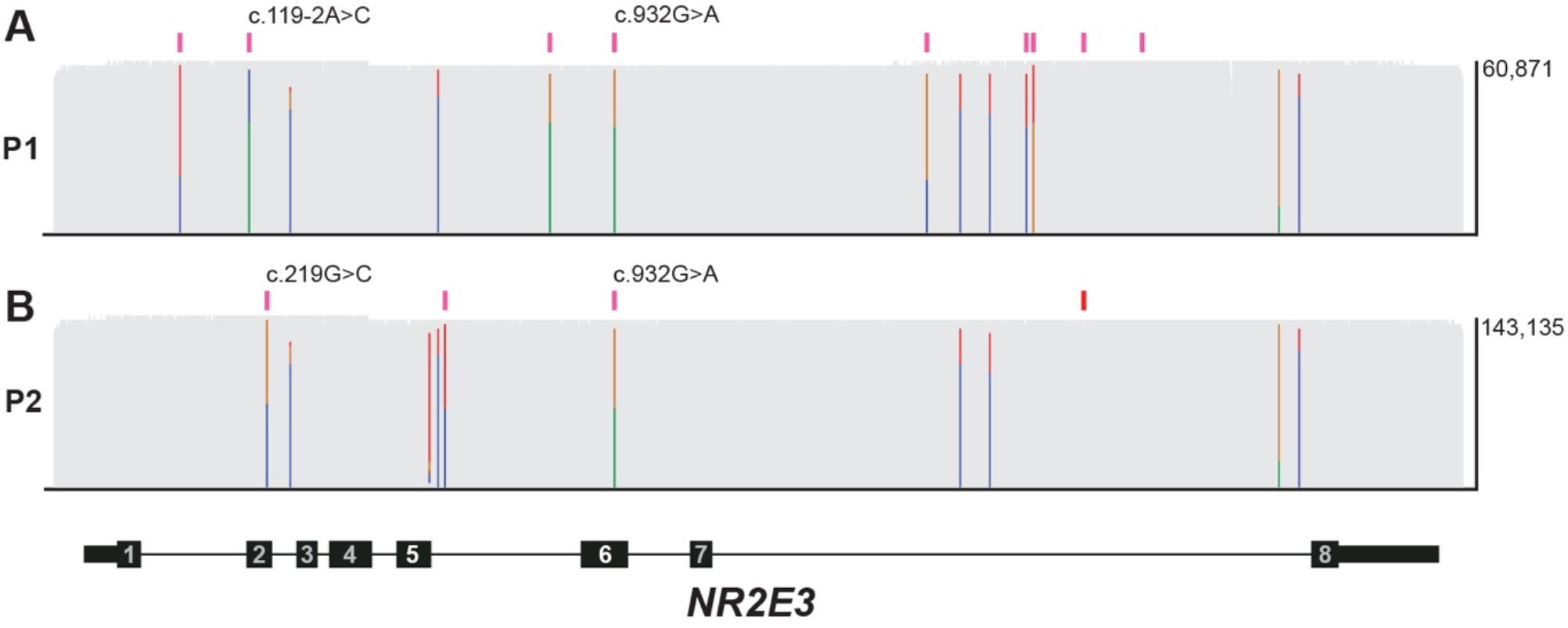
gDNA sequencing of the NR2E3 locus of patients carrying c.932G>A. **A)** An amplicon spanning the entire locus of *NR2E3* was sequenced using long reads. Variants are shown in pink (heterozygous) or red (homozygous) bars. Known pathogenic variants are labeled. **B)** As in **A**, the locus of *NR2E3* from patient P2 is shown.

