## Supplemental Figure 1 for "Exonic splice variant discovery using in vitro models of inherited retinal disease"

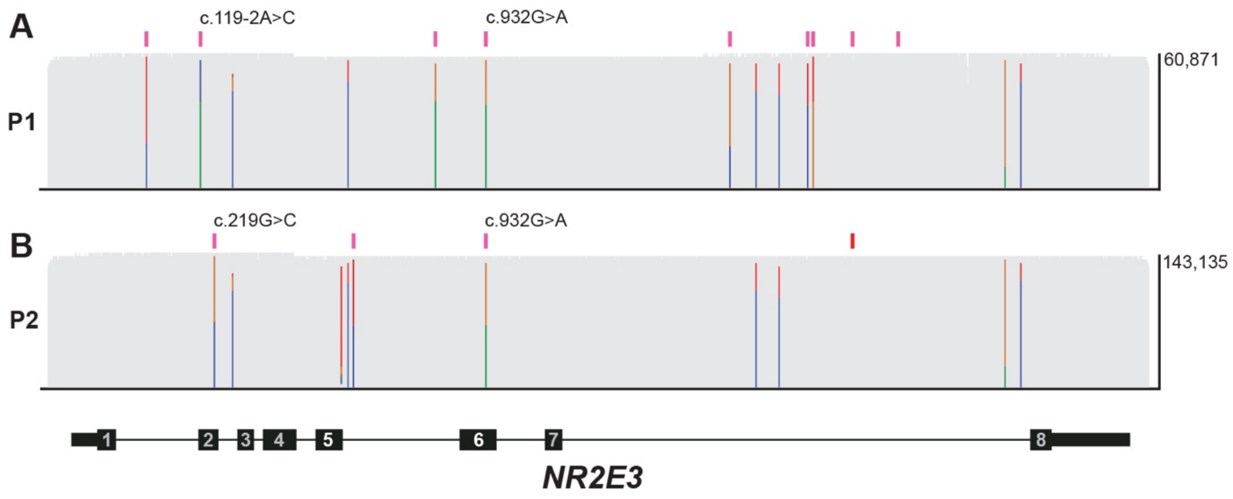


**Figure S1 gDNA sequencing of the NR2E3 locus of patients carrying c.932G>A. A)** An amplicon spanning the entire locus of *NR2E3* was sequenced using long reads. Variants are shown in pink (heterozygous) or red (homozygous) bars. Known pathogenic variants are labeled. **B)** As in **A**, the locus of *NR2E3* from patient P2 is shown.
